# Morphological, molecular, and functional characterization of mouse glutamatergic myenteric neurons

**DOI:** 10.1101/2023.09.18.558146

**Authors:** Jia Liu, Shaopeng Zhang, Sharareh Emadi, Tiantian Guo, Longtu Chen, Bin Feng

**Affiliations:** Department of Biomedical Engineering, University of Connecticut, CT 06269

**Keywords:** Glutamate, Vesicular glutamate transporter, myenteric neuron, mechanosensitive, calcium imaging

## Abstract

The enteric nervous system (ENS) functions largely independently of the central nervous system (CNS). Correspondingly, glutamate, the dominant neurotransmitter in the CNS and sensory afferents, is not a primary neurotransmitter in the ENS. Only a fraction (approximately 2%) of myenteric neurons in the mouse distal colon and rectum (colorectum) are positive for vesicular glutamate transporter type 2 (VGLUT2), the structure and function of which remain undetermined. Here, we systematically characterized VGLUT2-positive enteric neurons (VGLUT2-ENs) through sparse labeling with adeno-associated virus, single-cell mRNA sequencing (scRNA-seq), and GCaMP6f calcium imaging. Our results reveal that the majority of VGLUT2-ENs (29 out of 31, 93.5%) exhibited Dogiel type I morphology with a single aborally projecting axon; most axons (26 out of 29, 89.7%) are between 4 and 10 mm long, each traversing 19 to 34 myenteric ganglia. These anatomical features exclude the VGLUT2-ENs from being intrinsic primary afferent or motor neurons. The scRNA-seq conducted on 52 VGLUT2-ENs suggests different expression profiles from conventional descending interneurons. Ex vivo GCaMP6f recordings from flattened colorectum indicate that almost all VGLUT2-EN (181 out of 215, 84.2%) are indirectly activated by colorectal stretch via nicotinic cholinergic neural transmission. In conclusion, VGLUT2-ENs are a functionally unique group of enteric neurons with single aborally projecting long axons that traverse multiple myenteric ganglia and are activated indirectly by colorectal mechanical stretch. This knowledge will provide a solid foundation for subsequent studies on the potential interactions of VGLUT2-EN with extrinsic colorectal afferents via glutamatergic neurotransmission.

**New & Noteworthy:** We reveal that VGLUT2-positive enteric neurons (EN), although constituting a small fraction of total EN, are homogeneously expressed in the myenteric ganglia, with a slight concentration at the intermediate region between the colon and rectum. This concentration coincides with the entry zone of extrinsic afferents into the colorectum. Given that VGLUT2-ENs are indirectly activated by colorectal mechanical stretch, they are likely to participate in visceral nociception through glutamatergic neural transmission with extrinsic afferents.

## Introduction

The enteric nervous system (ENS) regulates gut mechanical motility, i.e., peristalsis and segmentation in a highly autonomous way and has been considered as a “second brain”. The ENS operates autonomously as evidenced by the presence of peristalsis, colonic migrating motor complex, and colonic transit in excised and isolated colon segments (37). The intricate neural circuitry that regulates the above motor movements consists of three major neural categories with distinct functions, including the intrinsic sensory neurons (intrinsic primary afferent neurons [IPANs]), interneurons, and motor neurons. The IPANs are the first neurons in the intrinsic reflexes that influence the patterns of motility, water and electrolyte secretion, and local blood flow by forming synaptic transmission to interneurons, motor neurons and other IPANs. The primary neurotransmitters for IPAN are acetylcholine, calcitonin gene-related peptide (CGRP), and tachykinin (13). Interneurons primarily locate in the myenteric plexus and can be categorized into two types, ascending and descending interneurons that project to excitatory and inhibitory motor neurons, respectively. Interneurons also project to other interneurons of the same type. Acetylcholine is the primary neurotransmitter for interneurons (36). Motor neurons innervate smooth muscles in the circular and longitudinal muscle layers, arterioles, and epithelium. The excitatory motor neurons predominantly use acetylcholine as their neurotransmitter with a small component of tachykinins (6). The inhibitory motor neurons primarily use nitric oxide as the neurotransmitter, with vasoactive intestinal peptide (VIP), adenosine triphosphate (ATP), and carbon monoxide (CO) as secondary neurotransmitters (5).

In contrast to the dominant role in extrinsic afferents, glutamate is considered a nonessential neurotransmitter in the ENS with only minor roles compared to the neurotransmitters summarized above. In fact, intracellular electrophysiological recordings and GCaMP6f calcium imaging studies of myenteric neural somata indicate that only a small fraction of myenteric neurons (∼10%) respond to glutamate application with a slow excitatory post synaptic potential (EPSP), which appears to be exclusively driven by the activation of excitatory metabotropic glutamate receptors (mGluR) (30, 39). Pharmacological studies with focal electrical stimulation confirm that synaptic transmission between glutamatergic nerve terminals and myenteric neurons are through excitatory mGluR (39). Further, fast EPSPs from activation of ionotropic glutamate receptors (iGluR) is absent in myenteric neurons (30), and glutamatergic receptor agonists L-Glu and ACPD do not alter the resting membrane potential of myenteric neurons (41). Finally, myenteric neurons are unresponsive to antidromic electrical stimulation of extrinsic afferents (35), which express vesicular glutamate transporters and presumably release glutamate into peripheral tissues (29). In sum, there is limited role of glutamate in the enteric neural circuitry despite being the major neurotransmitter for extrinsic afferents whose endings reside in the intestinal wall.

Using antibody staining and in situ hybridization, a prior report indicates that a small proportion of myenteric neural somata in the ENS of mouse colon express VGLUT2 (4), suggestive of their role in presynaptic glutamate release. In support, a recent study in mouse myenteric ganglia reveals abundant immunohistochemical presence of the presynaptic marker synaptophysin in VGLUT2-positive varicosities within the ganglia (39). We recently showed that VGLUT2-Cre labels a fraction of (∼2%) neurons in the myenteric ganglia (i.e., the VGLUT2-EN). Given that clinical evidence suggests a correlation between functional gastrointestinal disorders and altered glutamate signaling in the gastrointestinal tract (18), the VGLUT2-positive enteric neurons, although being the minority, could play critical roles in visceral nociception through glutamatergic neurotransmission to extrinsic sensory afferents. In this study, we implemented adeno-associated virus (AAV)-mediated sparse-labeling and single-cell mRNA sequencing (scRNA-seq) to uncover the morphology and molecular profile of VGLUT2-ENs. We also conducted functional studies by recording GCaMP6f signals from individual VGLUT2-ENs in intact colorectum while delivering mechanical colorectal stretch. Combining the morphological, molecular, and functional evidence, we for the first time delineate the VGLUT2-EN as a new functionally distinct group of enteric neurons.

## Materials and Methods

All experiments were reviewed and approved by the University of Connecticut Institutional Animal Care and Use Committee. All the mice used in the following experiments were housed in pathogen-free facilities which are Public Health Service assured and American Association for Accreditation of Laboratory Animal Care accredited following the Guide for the Care and Use of Laboratory Animals Eighth Edition. Mice resided in individual ventilated caging systems in polycarbonate cages (Animal Care System M.I.C.E.) and were provided with contact bedding (Envigo T7990 B.G. Irradiated Teklad Sani-Chips). Mice were fed ad lib with either 2918 Irradiated Teklad Global 18% Rodent Diet or 7904 Irradiated S2335 Mouse Breeder Diet supplied by Envigo and supplied with reverse osmosis water chlorinated to 2 ppm using a water bottle. Nestlets and huts were supplied for enrichment. Rodent housing temperature was set for 73.5 °F with a range from 70 to 77 °F. Humidity was set at 50% with a range of 35% to 65%. Mice were housed with a maximum of 5 animals per cage. All animals were housed on a 12:12 light-dark cycle. Animals were observed daily by the animal care services staff. Cages were changed every 2 weeks.

### Transgenic Mice

Transgenic mice of both sexes aged 8 -14 weeks were used for the study. Mice as young as 6.5 weeks of age were used for colonic AAV injection, but they will not be used in experiments until they reach 8 weeks of age. To identify the VGLUT2-ENs for morphological study and scRNA-seq, homozygous VGLUT2-Cre mice (strain no. 28863, The Jackson Laboratory, CT) were crossbred with homozygous floxed tdTomato reporter mice (Ai14, strain no. 7914, The Jackson Laboratory, CT) as reported previously (15). In offspring with heterozygous expression of Cre and floxed tdTomato, the VGLUT2 promotor drives the expression of the red fluorescent reporter tdTomato which fills the cytoplasmic space and large processes of VGLUT2-ENs near the neural somata.

For calcium imaging studies, the Ai95 mice carrying homozygous floxed GCaMP6f gene (strain# 28865, The Jackson Laboratory, CT) and homozygous VGLUT2-Cre mice (strain# 28863, Jackson Laboratory, CT) were crossbred. Offspring with heterozygous expression of Cre and floxed GCaMP6f were used for the experiments, in which VGLUT2-ENs express GCaMP6f, a fluorescent Ca2+ indicator for measuring changes in cytoplasmic free calcium as a metric of neural activities.

### Adeno-associated viral (AAV) vector for sparse labeling of VGLUT2-ENs

We implemented Cre-dependent AAV for sparse labeling of individual VGLUT2-ENs in VGLUT2-Cre mice (strain no. 28863, The Jackson Laboratory, CT) using AAV1-floxed-ChR-EYFP (#20298-AAV1, Addgene). From our preliminary study (data not shown) for assessing the viral transfection efficiency of different serotypes, we found that AAV9 transfects VGLUT2-ENs in almost all myenteric ganglia while AAV1 only transfects a fraction of the myenteric ganglia (∼20%). Since there are only 1 to 3 VGLUT2-ENs in each myenteric ganglia, Cre-dependent AAV1 enables sparse labeling of VGLUT2-ENs in VGLUT2-Cre mice to allow anatomic tracing of individual VGLUT2-ENs without any interference from other labeled neurons. We implemented the EYFP fused with Channelrhodopsin2 (ChR-EYFP) as the fluorescent reporter for our anatomic tracing, which is a membrane-bound marker and has been shown to transport more than 10 mm from the somata to peripheral nerve endings based on prior optogenetic studies on the dorsal root ganglia (12, 42) and neural tracing studies on myenteric neurons (16).

Mice were anesthetized by inhalation of 1.5% isoflurane, the distal colon was exposed by laparotomy, and the colon wall was injected with AAV1-floxed-ChR-EYFP (at 4-5 locations, ∼2-4 mm apart, 0.1 μL per site, titer >1x10^13^ vg/mL) by a syringe with a 33-gauge needle (model 7643-01, Hamilton Company, Arlington, MA). Any leakage from injection sites was immediately removed using a cotton-tipped applicator. After injection, the peritoneal cavity was rinsed with sterile saline. The muscle and skin were then sutured subsequently using absorbable polyglactin suture (VCP416H, Ethicon) and non-absorbable polypropylene suture (8698G, Ethicon) separately. To alleviate post-surgical inflammation and pain, meloxicam (2 mg/kg, Boehringer Ingelheim Vetmedica, Duluth, GA) was given three times in 72 hours for postoperative analgesia. After injection, the mice were allowed 20 days for sufficient AAV transfection.

### Colorectal tissue preparation and microscopic imaging for morphological study

Mice were deeply anesthetized and euthanized by transcardiac perfusion from the left ventricle with ice-cold oxygenated Krebs solution (in mM: 117.9 NaCl, 4.7 KCl, 25 NaHCO_3_, 1.3 NaH2PO_4_, 1.2 MgSO_4_, 2.5 CaCl_2_, and 11.1 D-glucose). The excised ∼30 mm of distal colorectum was removed from the mouse and immediately placed in ice-cold oxygenated Krebs solution. The colorectum was then cut along the mesentery, slightly stretched axially and circumferentially, and pinned flat in the Sylgard-lined Petri dish, and fixed with 10% buffered formalin at 4 °C overnight. The fixed tissue was washed with phosphate-buffered saline (PBS) three times (10 min each) to remove the fixative. Some tissues were incubated with a mouse biotin-conjugated monoclonal antibody against HuC/HuD (1:200, Fisher Scientific, East Greenwich, RI) for 48 h at 4 °C, and then incubated in Streptavidin conjugated with Alexa Fluor 488 (1:200, Jackson Immuno-Research, West Grove, PA) for 2 h at room temperature. Afterwards, the fixed or immune-stained colorectum was immersed in reflective index matching solution (RIMS, see Ref. (20) for details) for 30 min before being mounted in RIMS for subsequent microscopic imaging.

For VGLUT2-ENs distribution analysis, colorectum slides were imaged by Olympus BX51WI (Olympus, Waltham, MA) using a 4X objective lens (UPlanFLN 4X, Olympus, JP). The microscopic images were stitched and cells with tdTomato fluorescence were counted using 2D-stitching (27) and TrackMate (8) plugins respectively in ImageJ v1.5.3 (33). For the axonal projection study, the AAV-labeled tissue was observed by Zeiss LSM 510 Multiphoton confocal microscope using a 20X water-immersed objective (20X/0.75 HC PL APO, Leica). Images were taken as 2 μm-stepped z-stacks which covered the thickness of the projection of each traced axon. For each labeled enteric neuron, at least 10 adjacent 580x580 μm views were taken to ensure the coverage of at least 5 mm length of the axon for neurons with long processes. The confocal z-stacks were manually stitched using the 3D-stitching plugin (27), and the projection of individual axons was traced using the Simple Neurite Tracer (SNT) (1) plugin in ImageJ v1.5.3(33). To identify the cell size and dendrite shape, high resolution images of the somata of each AAV-labeled VGLUT2-ENs were taken by a 40x oil objective (40x/1.30 Plan Fluor, Nikon) using a Nikon A1R confocal microscope with the z-step set as 0.5 μm.

### Calcium imaging of colorectal myenteric plexus under electrical or mechanical stimulation

The excised ∼30 mm of distal colorectum was harvested from the VGLUT2-GCaMP6f mouse following the same euthanasia and perfusion protocol as described earlier. The colorectal section was dissected in ice-cold oxygenated Krebs solution and cut longitudinally along the mesenteric border. With serosal side up, one side of the mesenteric borders was pinned firmly onto a silicone elastomer substrate (Sylgard 184, Dow, MI), and the other side was hooked to a custom-built rake (1-mm interval, 22 mm long) to allow uniform circumferential stretch as reported previously (9, 10). The tissue was constantly perfused with 32–34 °C Krebs solution bubbled with carbogen (95% O_2_, 5% CO_2_). Nifedipine (1 μM) was added to the Krebs solution to inhibit spontaneous smooth muscle contraction. Electrical stimulation was delivered by a stimulus isolator (Model 701C, Aurora Scientific, Ontario, CA) via a round-tipped concentric electrode (external Φ0.55 mm, internal Φ0.125 mm; FHC, Bowdoin, ME), which was placed perpendicular to the serosal surface. Circumferential stretch was produced by a servo-controlled force actuator (Aurora Scientific, Aurora, Ontario, Canada). The delivery of electrical and mechanical stimuli was controlled by the Spike2 software (v7, Cambridge Electronic Design [CED], Cambridge, UK) running on an analog/digital interface (Power1401-3A, CED, Cambridge, UK).

At each region of interest, GCaMP6f signals were recorded using an upright microscope platform (BX51WI, Olympus, Waltham, MA) with a water immersion 10X objective (UMPLFLN 10XW, 0.3 NA). GCaMP6f signals were evoked by a fluorescent illuminator (Lumen 200, Prior Scientific, Cambridge, UK) through the 10X objective and recorded at 65 frames per sec using an ultra-low noise sCMOS camera (Xyla-4.2P, 82% quantum efficiency, Andor Technology, South Windsor, CT). We employed a segmentation algorithm ‘TrackMate’ in ImageJ (8) to determine the neuron location coordinates and fluorescence intensity of individual VGLUT2-ENs when the colorectum was subjected to electrical or mechanical stimulation. The signals were post-processed by MATLAB (MathWorks, Natick, MA) to eliminate the change in local background fluorescence caused by motion artifact especially during circumferential colorectal stretch. As shown in Fig. 1, we first electrically stimulated the VGLUT2-ENs by placing the concentrical electrode 300 μm away from the soma and delivering a 0.5 Hz stimulation (biphasic constant current stimulation, cathodic pulse first, 1.5 mA, 0.04 ms duration), an intensity sufficient to excite all neuronal tissues through the wall thickness based upon our prior study (10). We then waited for 5 min to allow VGLUT2-ENs to fully recover from the after hyperpolarization (AHP) before delivering mechanical circumferential stretch: 0 to 140% stretch ratio in 10 sec. The 140% stretch ratio usually evokes a maximum stretch force of 0.4 to 0.6 N to correlate with intraluminal colorectal distension of 80 to 100 mmHg assuming the colorectum as a thin-walled cylinder (9, 34). In each VGLUT2-EN, the response to circumferential stretch was recorded four times: two baseline responses separated by 20 min, 20 min after switching to bath solution containing 200 μM hexamethonium chloride (nicotinic cholinergic blocker, Sigma Aldrich), and 25 min after wash-off.

**Fig. 1.**
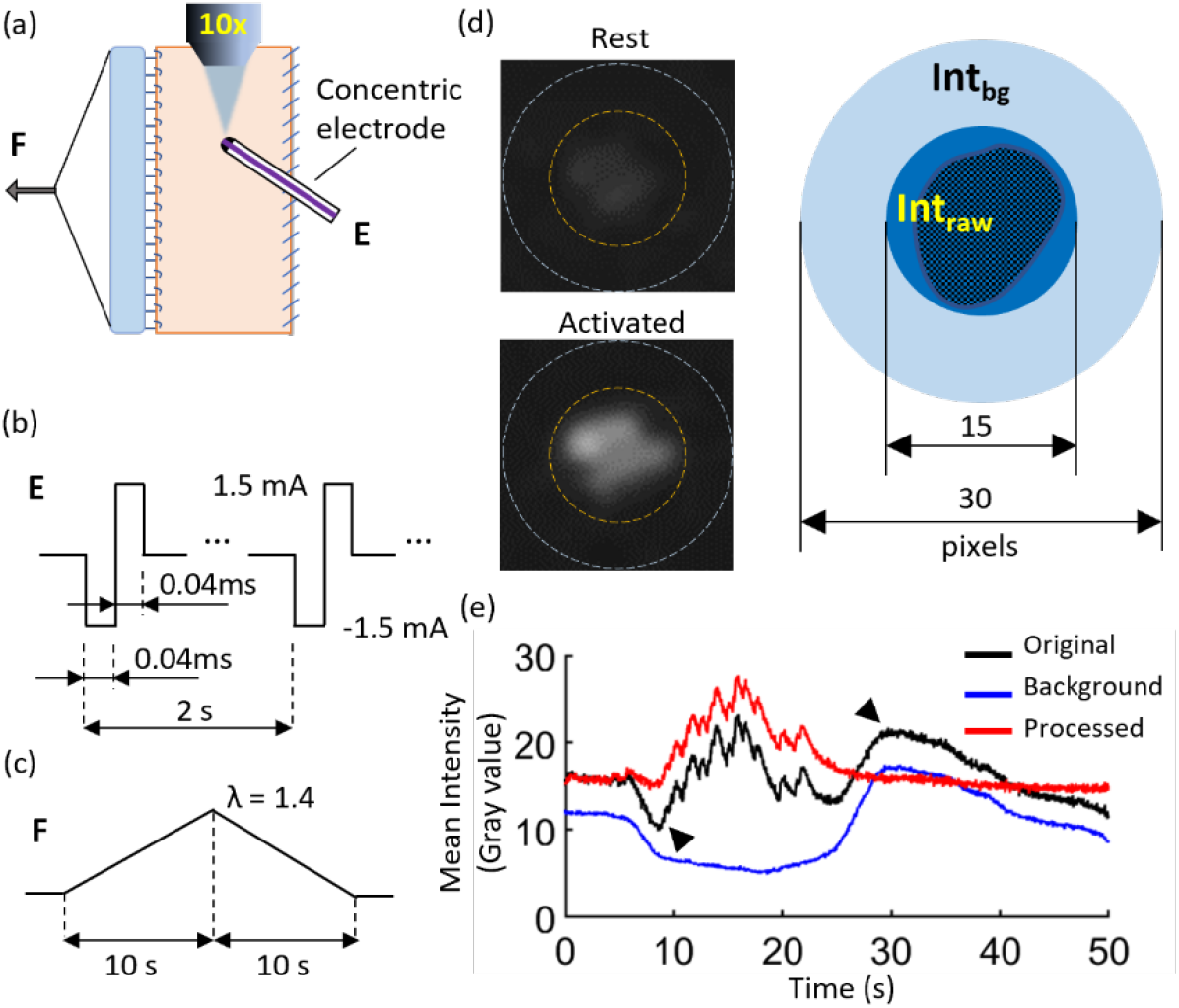
GCaMP6f recordings from VGLUT2-ENs in whole mouse colorectum (∼22 mm long) undergoing electrical and mechanical stimuli. (a) Schematic of the flattened colorectum with serosal side up for ex vivo electrical stimulation by a concentric electrode placed on the colorectal serosa and circumferential mechanical stretch via a custom-built rake (1 mm intervals, 22 mm wide). GCaMP6f signals were recorded from each VGLUT2-EN through an 10X objective at 65 frames per second. (b) Electrical stimulation consists of 0.5 Hz biphasic pulse trains of 0.04 ms duration and 1.5 mA amplitude. (c) The circumferential stretch is delivered as a slow ramp from zero to 140% stretch ratio in 10 sec. (d) Representative GCaMP6f recordings from a VGLUT2-EN in resting and activated states, respectively. (e) Signal processing by background subtraction to reveal GCaMP6f signals recorded from a VGLUT2-EN subjected to circumferential colorectal stretch.

A significant challenge from conducting GCaMP6f recordings from individual VGLUT2-ENs of ∼20 μm in size is the significant motion artifact from large strain colorectal deformation (up to 140% stretch ratio), which causes varying background GCaMP6f signals during stretch. To address the adverse impact of variable background signals, we developed a post-processing algorithm to deduce the background signal from a fine ring-shaped region surrounding the individual VGLUT2-EN. As illustrated in Fig. 1d, we calculated the time-dependent average fluorescence intensity (*Int*_*raw*_ (*t*)) of a circular region that covered the VGLUT2-EN soma, and the circle center coordinates by the TrackMate plugin in ImageJ. Using the center coordinates, we developed a custom-built MATLAB program (v2023, MathWorks Inc.) to calculate the time-dependent average fluorescence intensity (*Int*_*bg*_(*t*)) of the surrounding ring with the outer diameter twice the diameter of the circle that covers the soma, which was assumed as the local background signal. Then, the background-corrected GCamp6f from the VGLUT2-EN soma, *Int*_*cel*_ (*t*), was calculated as,

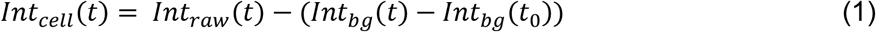

where *t*_0_ is the initial time point of each measurement. Before the background correction, there exists significant background variations in the fluorescent intensity as indicated by arrowheads in Figure 1e (black trace). After correction, the fluorescent intensities have a stable baseline to reveal evoked GCamp6f signals in the VGLUT2-EN soma (Fig. 1e, red trace).

### Single cell mRNA sequencing (scRNA-seq)

The excised ∼30 mm of distal colorectum was harvested from the VGLUT2-tdTomato mouse following the same euthanasia and perfusion protocol as described earlier. The colon section was dissected in ice-cold oxygenated Krebs solution to remove the anus and most of the mesentery and was flushed with Krebs solution to remove the fecal matter. The longitudinal-muscle-myenteric-plexus (LMMP) was gently peeled off from the colorectum using a wet cotton swab following a previously reported procedure (19). The isolated LMMP was trimmed into small pieces (∼4 x 4 mm) and enzymatically digested following a previously reported protocol with slight modification (40). Briefly, the tissue was collected in 15 ml conical tubes containing Hank’s balanced salt solution (HBSS, Sigma, Cat#: H6648-1L) and centrifuged at low speed (300 rpm, 5 minutes, room temperature). The supernatant was aspirated, and the tissue was resuspended in a pre-warmed collagenase digesting solution at 37°C, i.e., 2 mg/ml collagenase (type IV, Worthington Biochemical, Lakewood, NJ) in HBSS medium containing 0.5 mM CaCl_2_ and 10 mM HEPES. The tissue was incubated in the collagenase solution in a 37 °C water bath for 30 min with manual rotation and inverting every 5-10 minutes. Following centrifugation, the resulting pellet was washed twice with 10 ml pre-warmed HBSS. Next, the tissue was incubated with 1 ml pre-warmed (37 °C) trypsin solution (0.05% Trypsin-EDTA, Cat#: 25300-054, Gibco BLR) in a 37 °C water bath for 15 minutes. Following centrifugation, the pellet was triturated in DMEM/F12 culture media containing 10% FBS serum and 1% Penicillin-Streptomycin. Then, the cell suspension was pipetted onto precoated Laminin coverslips at 150-170 μl on each coverslip (Cat#: 72298-01, Electron Microscopy Science), placed in 35 mm petri dishes, and incubated for 3 hours in an incubator (37 °C, 5% CO_2_). Next, each petri dish was filled with 2 ml of warm DMEM-F12 culture media containing 10% FBS and 1% penicillin-streptomycin to submerge the coverslips and kept in the incubator overnight.

The dissociated neurons were cultured in the incubator (37 °C, 5% CO_2_) for 18 hours, a duration that is optimized for sufficient neural attachment to the coverslip to avoid cell loss. Also, the loose attachment from 18-hour culture enabled easy cell picking with a microcapillary. Right before the cell picking, the debris was removed by changing the culture medium and the 35-mm petri dish was mounted onto an upright fluorescent microscope (BX51WI, Olympus, Waltham, MA). Each VGLUT2-EN positive for tdTomato fluorescence was gently aspired into a heat-polished and autoclaved glass microcapillary (20 μm opening) by negative pressure generated via a high-precision manual pneumatic microinjector (CellTram 4r Air, Eppendorf, Hamburg, Germany). The VGLUT2-EN was then expelled into a single PCR tube containing 5 μL lysis mix (SeqWell, Beverly, MA) via positive pressure also generated by the microinjector. In some cell cultures, myenteric neurons were labeled by NeuroFluor™ NeuO, which is a green fluorescent probe specific for neuronal cells, and the cells that have green fluorescence but are negative for tdTomato fluorescence were retrieved as control group, i.e., the non-VGLUT2-ENs. The libraries for cDNA sequencing were prepared using a commercial scRNA-seq kit (PWSCR96, SeqWell, MA). The cDNA library was subjected to quality control by a QuantiFluor dsDNA system (E2671, Promega, WI) and an Agilent D5000 ScreenTape system before being sent to scRNA-seq (NovaSeq 6000, Illumina, San Diego, CA) at UConn Center for Genome Innovation. The read depth per cell is 5 million reads/cell and the read length is 100 bp.

### Data statistics

Statistical analysis was performed using SigmaStat. The student’s t-test was used to compare the average value between two groups. One-way ANOVA was applied to compare between multiple groups. Chi-square test was implemented for comparing proportions. Significance was set as p < 0.05 or specified otherwise. Results are expressed as mean ± SEM. At least three animals were used for each experimental condition. ‘n’ refers to the number of cells or number of tests as indicated, ‘N’ refers to the number of animals.

## Results

### Distribution of VGLUT2-ENs in mouse distal colorectum

To thoroughly quantify the distribution of the VGLUT2-ENs in mouse distal colorectum, tdTomato fluorescence signal over the entire VGLUT2/tdTomato colorectum was detected by imaging the entire distal 30 mm colorectum with a 4X objective (Fig. 2a) and stitching the images together. The location of each VGLUT2-EN in the colorectum was determined by automatically extracting the pixel coordinates of the center of the cell somata using ImageJ. Displayed in Fig. 2b is topological distribution of VGLUT2-EN identified from 3 colorectums shown as the heatmap of neurons per 0.64 mm^2^ grid. The quantified VGLUT2-EN neural density in Fig. 2C indicates that rectal region contains significantly fewer VGLUT2-ENs per area than the colonic and intermediate regions. By immunostaining the myenteric neural somata with anti-Hu antibody, we identified 6424 myenteric neurons from 62 field of views of 3 mouse colorectums, of which 1.8% (121 neurons) are VGLUT2-ENs (Fig. 3); all VGLUT2-ENs are positive for anti-Hu. The proportion of VGLUT2-ENs determined from the current study is consistent with the ∼2% VGLUT2-positive neurons in the ENS reported previously (3, 16). As shown in the bar graph in Fig. 3, the average proportion of VGLUT2-ENs in myenteric neurons is significantly higher in the intermediate region (3.07±0.38%) than in the colonic (1.56±0.37%) or rectal (1.77±0.46%) regions (One-way ANOVA, F = 4.19, p = 0.02).

**Figure 2.**
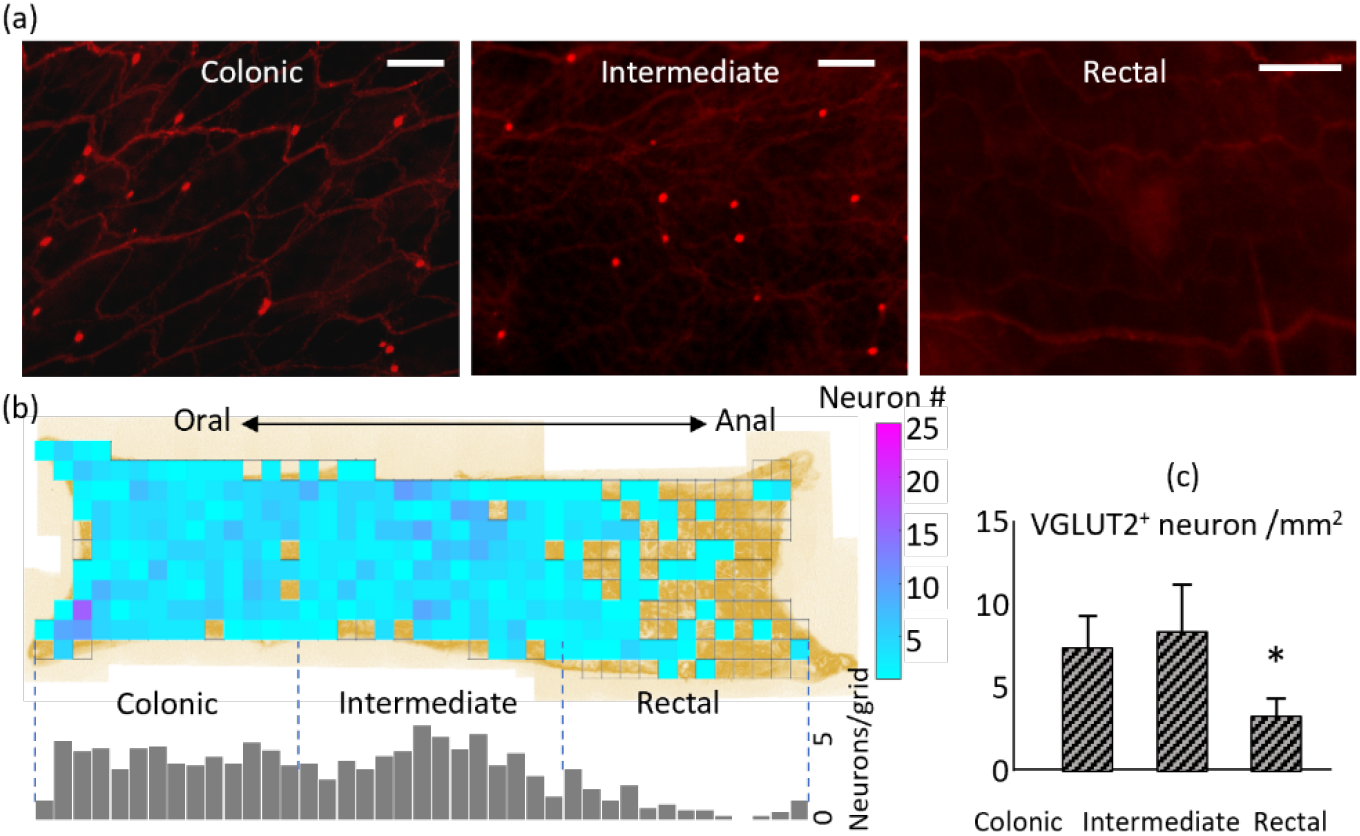
Area distribution of VGLUT2-ENs in mouse colorectum. (a) Representative fluorescent images of mouse colorectum captured by a 4X objective showing VGLUT2-ENs in red. (b) Heatmap and histogram plots showing the distribution of VGLUT2-ENs in a flattened mouse colorectum. (c) Area density of VGLUT2-ENs in mouse colorectum. The density at rectal region is significantly lower than at colonic and intermediate regions. Scale bar: 100 μm. *P<0.05.

**Fig. 3.**
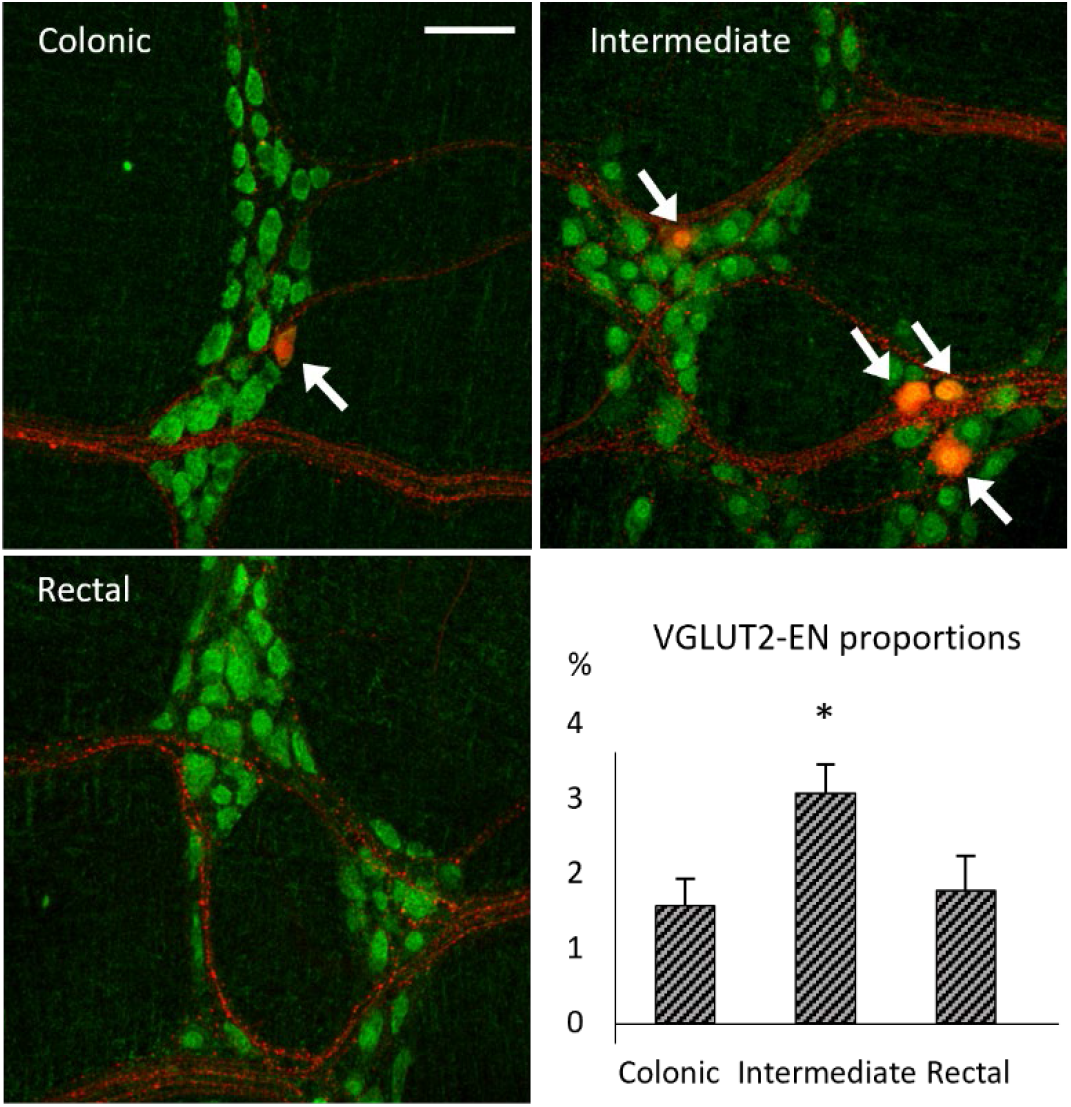
Proportions of VGLUT2-ENs in myenteric neurons. The colorectum from VGLUT2/tdTomato mice was whole mount stained by Anti-Hu antibody (green). The VGLUT2-ENs were labeled with red fluorescence (tdTomato). The proportion of VGLUT2-ENs in the myenteric neurons is significantly higher in the intermediate region than in the colonic and rectal regions. Scale bar: 50 μm.

### Morphological characterization of sparsely labeled VGLUT2-ENs using Cre-dependent AAV

To conduct morphological tracing of VGLUT2-ENs, we implemented sparse genetic labeling by injecting Cre-dependent AAV carrying the membrane-bound marker Channelrhodopsin2-EYFP (AAV1-floxed-ChR-EYFP) into the colon wall. Displayed in Fig 4a is a typical image of AAV-labeled colorectum from a VGLUT2/tdTomato mouse. In the field of view, only one VGLUT2-EN (green, yellow arrow) out of five (red) were labeled by the AAV; four VGLUT2-EN were not labeled (white arrows). The sparse AAV labeling coupled with the small proportion of VLUT2-EN in the myenteric ganglia (1.88%) enables anatomic tracing of individual VGLUT2-ENs of up to 10 mm long. We characterized the morphology of 31 sparse-labeled VGLUT2-ENs, tracing their axons towards the end in the whole-mount colorectum. Shown as a typical example in Fig. 4b, most traced VGLUT2-ENs (26/31) have Dogiel type I morphology (2) with a single long axon (>4 mm) projecting aborally to traverses 19 to 34 myenteric ganglia. They typically have small-sized somata (< 250 μm^2^) with lamellar dendrites showing tapering tips as in Fig. 4b inset. Very few of the VGLUT2-ENs (2/31) showed Dogiel type II morphology with multiple short axons (<3 mm) projecting circumferentially (Fig. 4c), whose somata are also small (<250 μm^2^) with lamellar dendrites showing enlarged tips (Fig. 4c inset).

**Figure 4.**
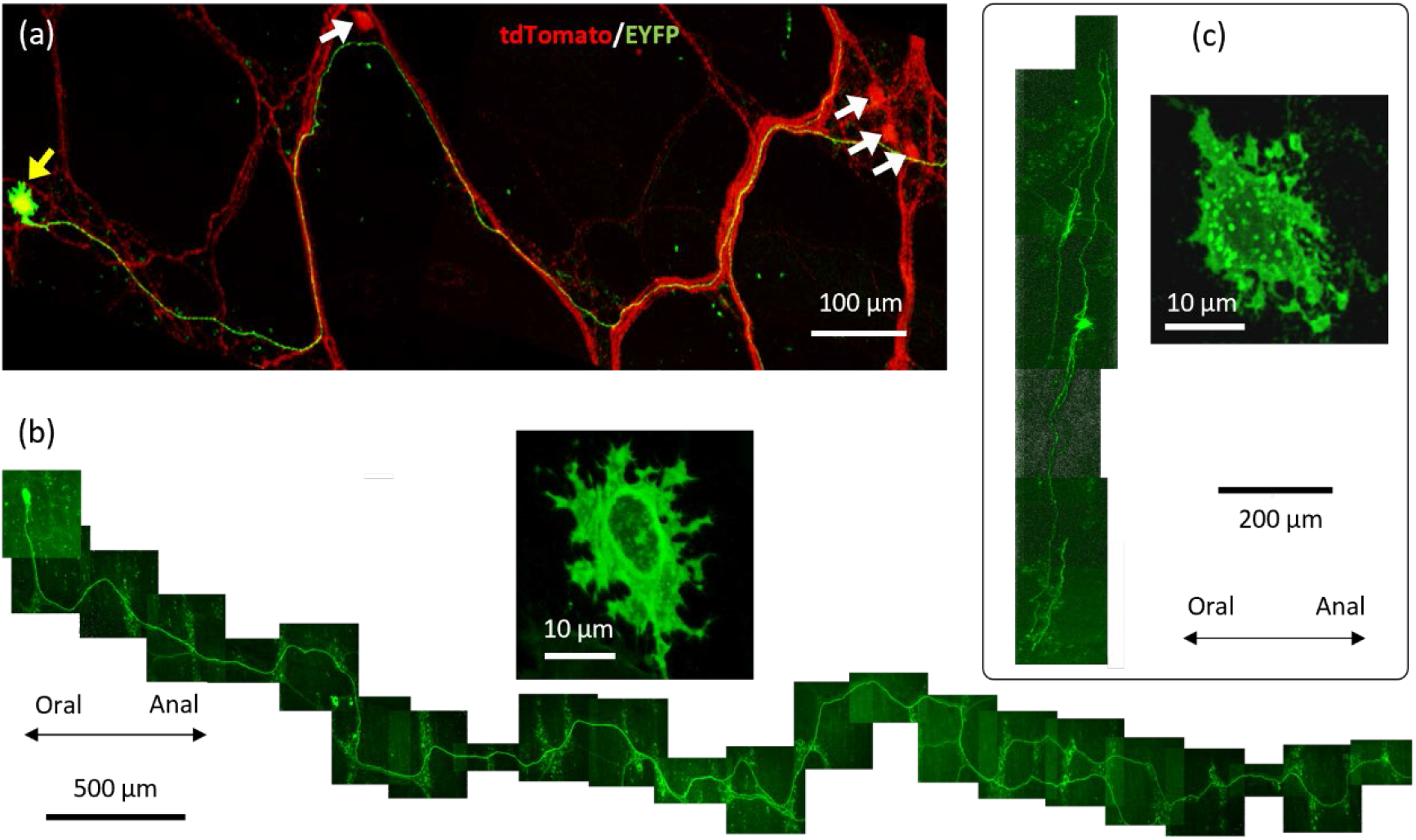
Morphological characterization of VGLUT2-ENs using sparse AAV tracing. (a) A z-stack confocal fluorescent image showing the successful sparse labeling of a VGLUT2-EN by AAV1-floxed-ChR-EYFP (yellow arrow). Four other VGLUT2-ENs in the field of view were not labeled (white arrow). (b) A representative VGLUT2-EN that has Dogiel type I morphology with a single aborally projecting axon of more than 4 mm. They typically have small-sized somata (< 250 μm^2^) with lamellar dendrites showing tapering tips (inset). (c) Very few VGLUT2-ENs have Dogiel type II morphology with multiple circumferentially projecting axons of less than 4 mm. Their somata are also small (<250 μm^2^) with lamellar dendrites showing enlarged tips (inset).

As summarized in Fig. 5a, the major category of VGLUT2-ENs (26/31, 83.9%) have Dogiel type I morphology with a single aborally projecting axon. The five outliers include two Dogiel type II neurons with multiple axons projecting circumferentially and three Dogiel type I neurons projecting either orally or circumferentially (26). Within the major category, 88% (23/26) have long projecting axons of more than 4 mm. In contrast, all five outliers have projecting axons no more than 4 mm. Most (23/31, 74.2%) VGLUT2-ENs have lamellar dendrites with tapering tips (Fig. 5 inset a1), and they typically fall into the major category. Five of the 31 VGLUT2-ENs have spiny dendrites (Fig. 5 inset a2) and they are all in the major category. Four of the 31 VGLUT2-ENs have lamellar dendrites with enlarged tips (Fig. 5 inset a3), which are all outliers. We also characterized the soma area of 75 sparse-labeled VGLUT2-ENs, including the 31 traced neurons and 44 neurons that cannot be traced due to interference from other labeled neurons. Displayed in Fig. 5b is the soma size distribution showing three groups by triple Gausian fit: small (192.44 ± 0.71, <260 μm^2^), medium (288.21 ± 0.92, 260-330 μm^2^), and large neurons (423.19 ± 13.88, >330 μm^2^). Most VGLUT2-ENs have small soma sizes (51/75, 68%), including the two Dogiel type II neurons in Fig. 5a.

**Fig. 5.**
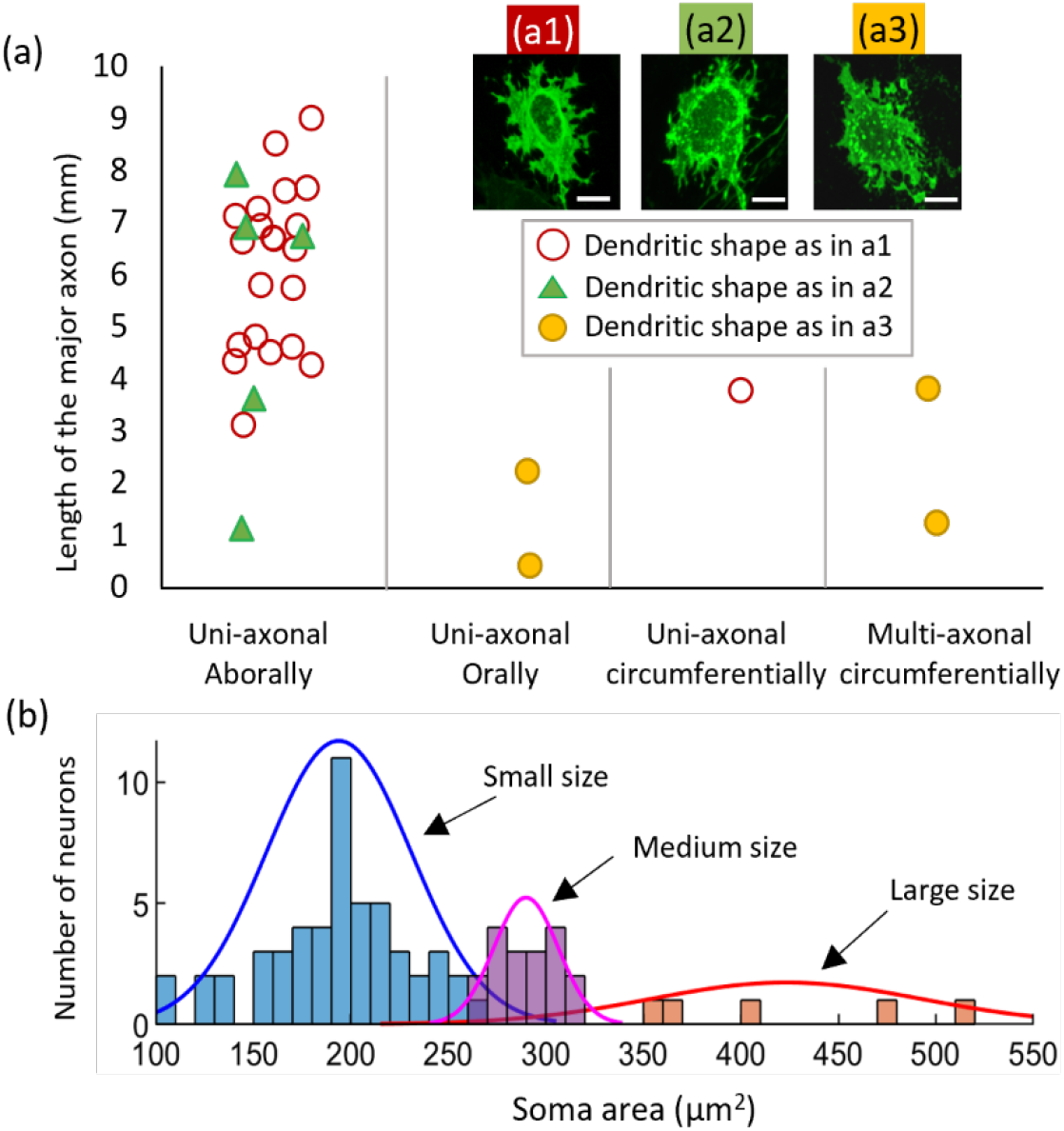
Morphological categorization of VGLUT2-ENs. (A) The major category of VGLUT2-ENs (25/31) have Dogiel type I morphology with a single aborally projecting axon. A small fraction of VGLUT2-ENs have Dogiel type II morphology with multiple axons projecting circumferentially. There are three exceptions of Dogiel type I VGLUT2-ENs with a single axon projecting orally or circumferentially. There are three major dendritic shapes as shown in inset a1, a2 and a3. (b) Soma size distribution and triple Gaussian fit of 75 sparse labeled VGLUT2-ENs.

### Molecular profiles of VGLUT2-ENs by scRNA-seq

As illustrated in Fig. 6a, 65 VGLUT2-ENs positive for red fluorescent signals were manually retrieved for scRNA-seq. As a control group, 33 myenteric neurons positive for the green neuronal marker (NeuroFluor) but negative for tdTomato (non-VGLUT2-ENs) were also harvested for scRNA-seq. Displayed in Fig. 6b are the expression level of VGLUT2 as quantified by read per kilobase per million (RPKM) in both VGLUT2-ENs and non-VGLUT2-ENs groups. All non-VGLUT2-ENs showed low express level of VGLUT2 with log(RPKM) below 0.01. In the VGLUT2-EN group, most neurons (61/65) showed significant level of VGLUT2 expression with log(RPKM) over 1. To remove non-neuronal cells from the pool of sequenced VGLUT2-EN, we extracted the expression levels of four neuronal markers NEFH, NEFL, MEFM, and PRPH and one astrocyte marker GFAP as shown in Fig. 6c. We removed 9 cells that either have high GFAP expression with log (RPKM) greater than 0.1 or low expression in all four neuronal markers (below the four dashed lines). Within the 52 screened VGLUT-ENs positive for neuronal markers, we conducted Principal Component Analysis (PCA) on the gene expression level of 15 well accepted and curated myenteric neuron markers (24, 28, 31, 32): 1) Calca, Calcb, Chat, Slc18a3, Nefm, Calb1, and Calb2 for IPAN; 2) Chat, Slc18a3, Calb2, and Tac1 for excitatory motor neuron; 3) Nos1, Vip, and Npy for inhibitory motor neuron; 4) Chat, Slc18a3, Nos1, and Vip for descending interneuron; 5) Chat, Slc8a3, Calb2, and Sst for ascending interneuron. Displayed in Fig. 6d is the clustering analysis results from PCA on the 15 gene markers, which indicate a dominant cluster of VGLUT2-ENs (43/52, 82.70%). By comparing the expression levels of the mechanosensitive ion channels Piezo1 and Piezo2 in the non-VGLUT2-

**Figure 6.**
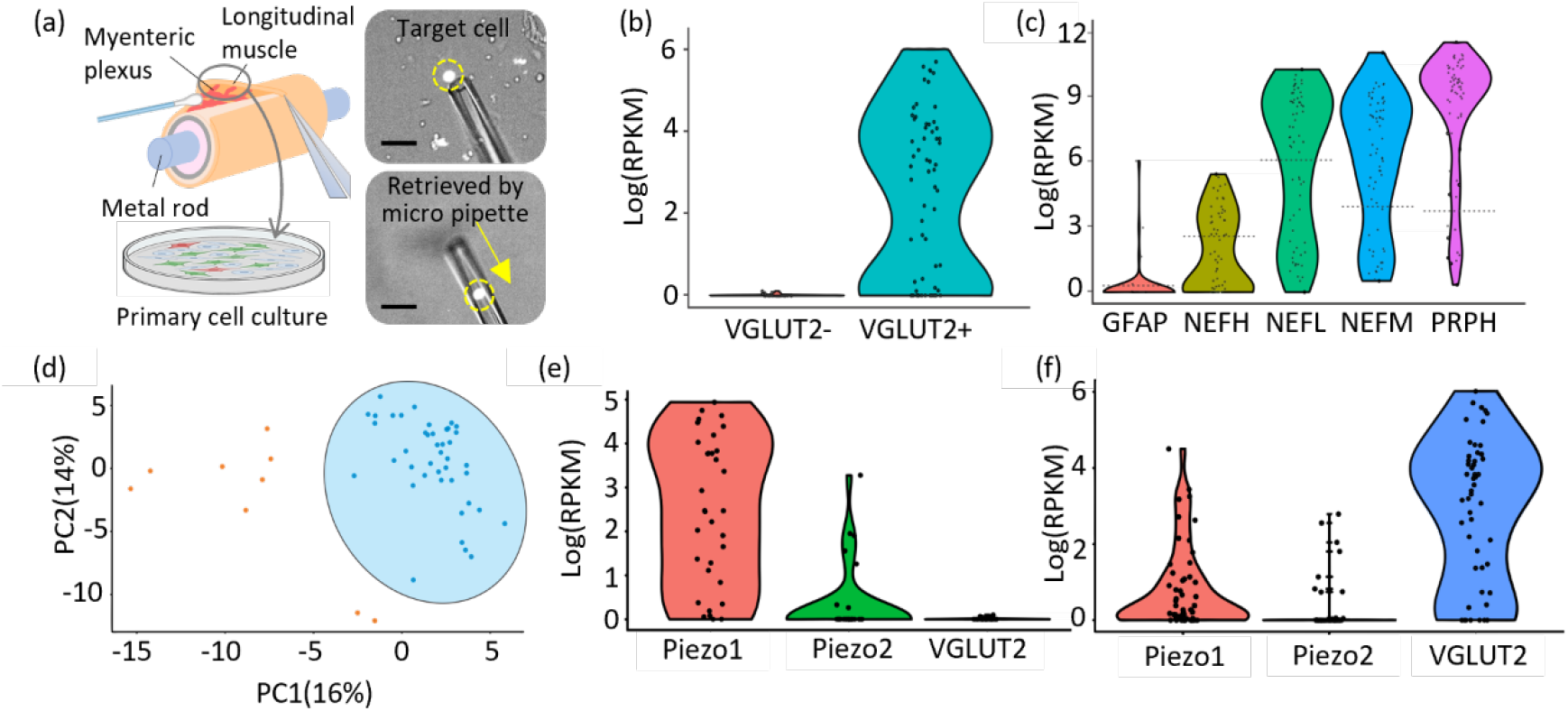
Characterize the molecular profiles of VGLUT2-ENs and non-VGLUT2-ENs by scRNA-seq. (a) Schematic and microscopic images showing the procedure of retrieving individual VGLUT2-ENs from dissociated longitudinal-muscle-myenteric-plexus (LMMP) harvested from VGLUT2/Ai14 colorectum. (b) The expression level of VGLUT2 gene in VGLUT2-EN and non-VGLUT2-EN as quantified by read per kilobase per million (RPKM). (c) Screening the VGLUT2-ENs with neuronal cell markers (NEFH, NEFL, NEFM, PRPH) and enteric glial cell marker (GFAP). (d) Principal component analysis to cluster VGLUT2-ENs. (e) Expression level of mechanosensitive ion channels piezo1 and piezo2 in non-VGLUT2-ENs. (f) Expression level of piezo1 and piezo2 in VGLUT2-ENs.

ENs (Fig. 6e) and VGLUT2-ENs (Fig. 6f), we find significantly less expression level of piezo1 and piezo2 genes in the VGLUT2-EN population as compared with myenteric neurons negative for VGLUT2.

In addition, the anatomical findings in Figs. 4 and 5 exclude VGLUT2-EN from being either IPAN or motor neurons, suggesting that they are descending inhibitory neurons. In Figure 7, we present the expression levels of published markers for descending interneurons in VGLUT2-ENs, including calcium-binding proteins (calb1, calb2), CGRP (calca1, calca2), nitric oxide synthases (nos1), vasoactive intestinal peptide (VIP), and serotonin synthesis/reuptake (dcc, slc6a4) (36). When compared with the expression levels of molecules involved in synaptic glutamatergic (slc17a6) and cholinergic transmission (chat and slc18a3c), it is evident that the expression levels of these descending interneuron markers are relatively low.

**Figure 7.**
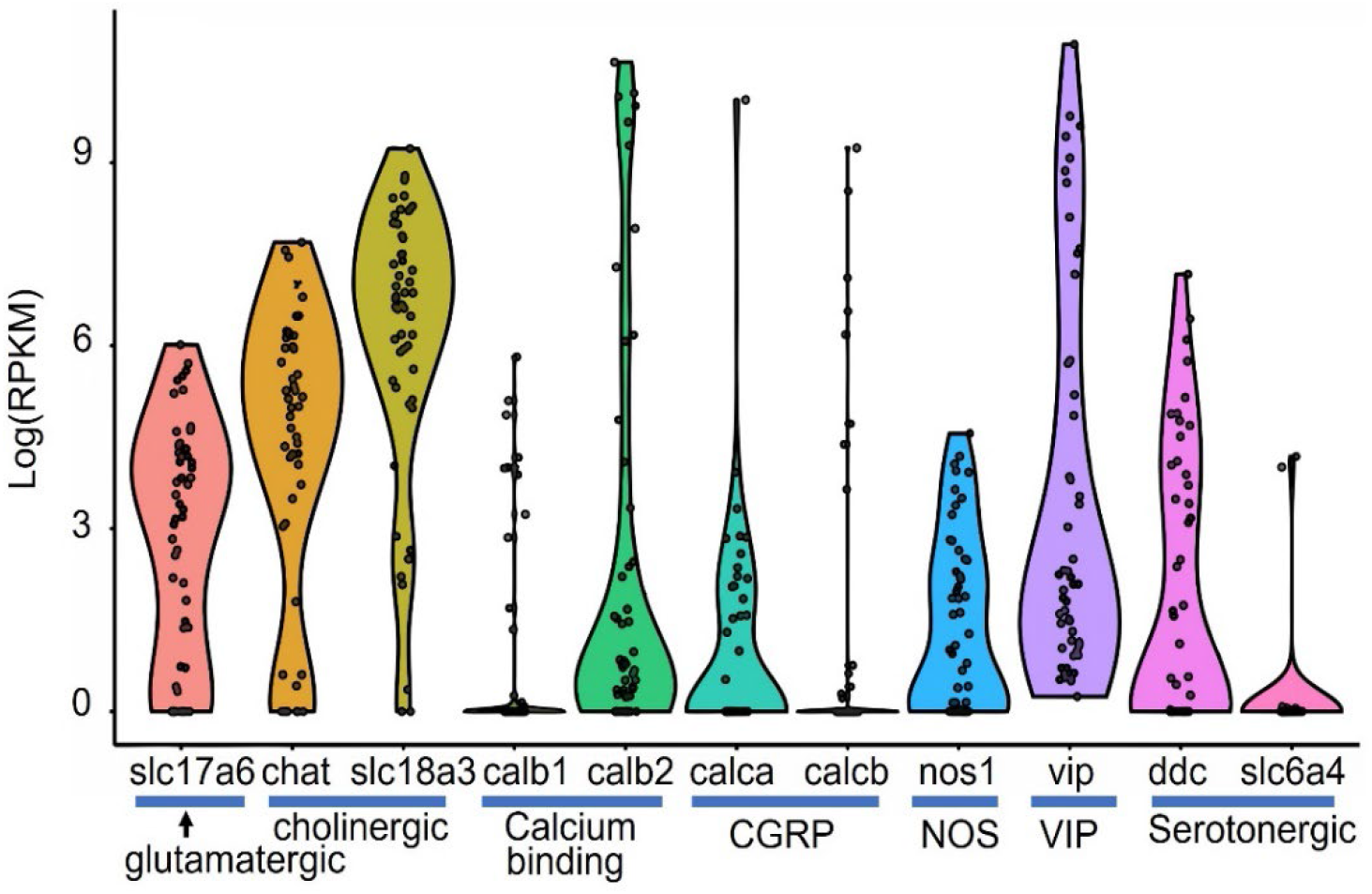
The expression levels of molecular markers for descending interneurons in VGLUT2-ENs, as compared to the levels of molecules involved in synaptic glutamatergic and cholinergic transmissions.

### Functional characterization of VGLUT2-EN by electrical and mechanical colorectal stimuli

Electrical stimulation delivered by the concentric electrode induced transients of GCaMP6f signals in VGLUT2-ENs that follow the stimulus frequency of 0.5 Hz (Fig. 8a). The recorded GCaMP6f from a typical VGLUT2-EN undergoing colorectal stretch was stacked and shown in Fig. 8b, from which the evoked GCaMP6f transients were derived by tracking the moving VGLUT2-EN and conducting background subtraction (c.f. Fig. 1). Displayed in Fig. 8c are the representative responses of a VGLUT2-EN to colorectal stretch before (control) and 20 min after switching the bath solution to containing hexamethonium (hexam), a potent inhibitor of the nicotinic acetylcholine receptor. The response to stretch was also recorded 25 min after washing off the hexamethonium. We identified a total of 215 VGLUT2-ENs with positive responses to electrical stimulation. Among them, 181 VGLUT22-ENs (84.2%) responded to circumferential colorectal stretch. The pooled results in Fig. 8d show that hexamethonium significantly inhibited the responses to colorectal stretch, which was recovered after wash (one-way ANOVA, F = 48.63, p < 0.001; post-hoc comparison, hexam vs control 2, p<0.001; hexam vs. wash, p<0.001).

**Figure 8.**
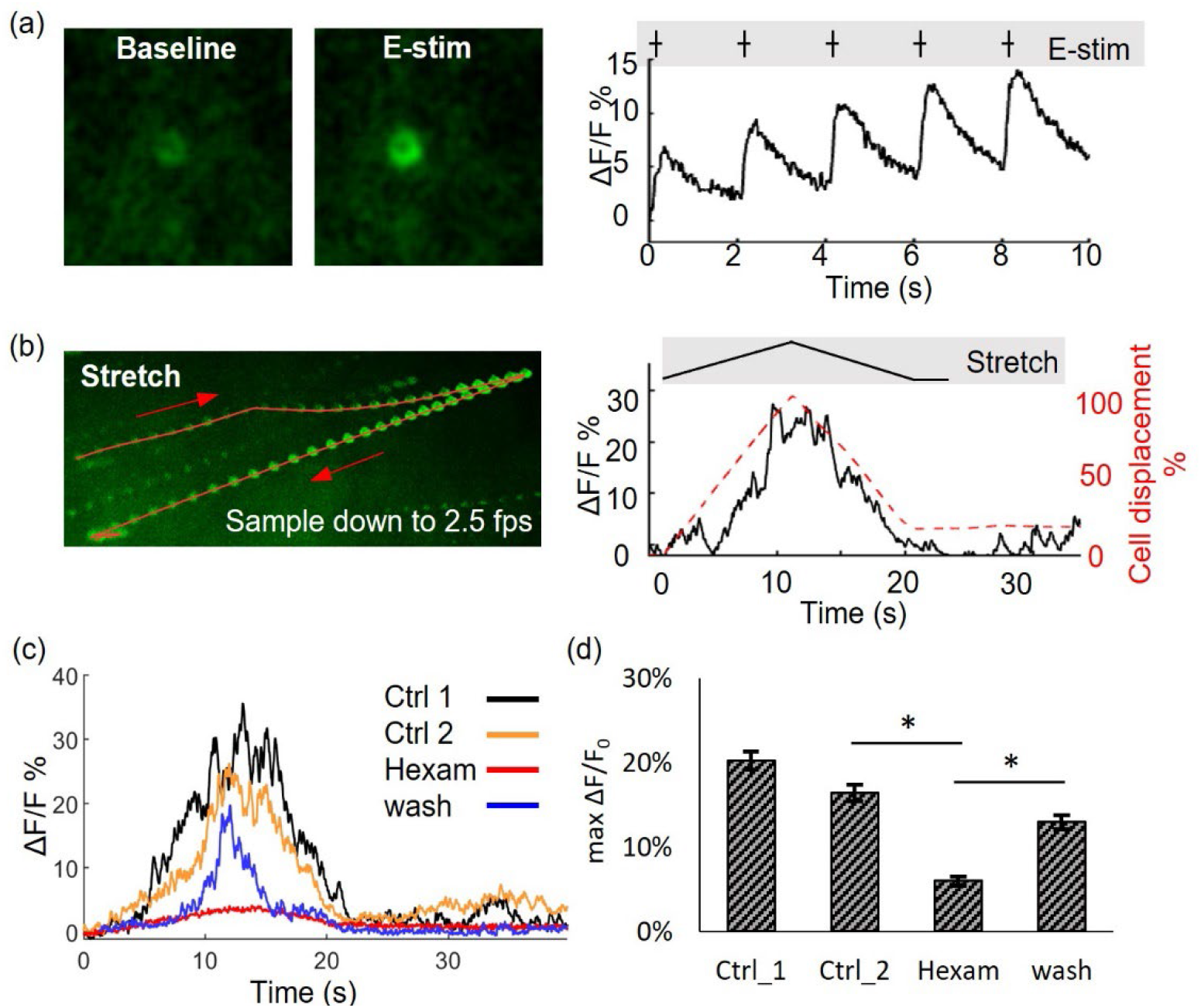
Functional characterization of VGLUT2-EN to mechanical colorectal stretch by GCaMP6f recordings. (a) Representative responses of a VGLUT2-EN to electrical stimulation and circumferential colorectal stretch were shown in (a) and (b), respectively. (c) Typical GCaMP6f responses of an VGLUT2-EN to colorectal stretch before and after bath application of hexamethonium (hexam, 200 μM). (d) In 181 tested VGLUT2-EN, hexamethonium significantly inhibited the GCaMP6f responses to stretch, which were partially recovered after wash.

## Discussion

In our current study, we conducted a comprehensive investigation into the distribution of VGLUT2-positive enteric neurons (VGLUT2-ENs) in the distal colorectum. Instead of random microscopic sampling, we meticulously examined the entire flattened tissue using the VGLUT2-Cre promoter. Our analysis confirms the specificity and efficiency of VGLUT2-Cre in labeling VGLUT2-ENs, as evidenced by single-cell RNA sequencing data, which demonstrate a significant expression of VGLUT2 in most, if not all VGLUT2-ENs but the absence of VGLUT2 expression in almost all non-VGLUT2-ENs. It is important to note that all VGLUT2-EN somata were detected in the myenteric plexus and were absent in the submucosal plexus, consistent with our prior research in mice (16). However, we acknowledge the possibility of VGLUT2-positive enteric neurons in the submucosal plexus in other species, particularly based on observations of VGLUT2-like immunoreactivity in the submucosal layer of human colon biopsies from our unpublished data. In this study, our focus is exclusively on VGLUT2-positive neurons in the myenteric plexus.

Despite VGLUT2-ENs constituting a fraction (1.88%) of myenteric neurons based upon the current and a prior study (16), they are evenly distributed throughout the colorectum, present in nearly every myenteric ganglion circumferentially. Along the longitudinal axis, VGLUT2-EN density remains fairly consistent in the colonic region, with a slight decrease toward the rectal region due to the reduced density of total myenteric neurons in the rectum. Notably, we observed a higher proportion of VGLUT2-ENs in the intermediate region between the colon and rectum, which coincides with the entry point of extrinsic afferent innervations from both lumbar splanchnic and pelvic nerve pathways (11). Extrinsic colorectal afferents prominently express all subtypes of ionotropic and metabotropic glutamate receptors from a scRNA-seq study on retrograde labeled neural somata in the DRG (17). The coexistence of VGLUT2-ENs and colorectal afferent endings in the intermediate region between the colon and rectum strongly suggests the potential for glutamatergic neural transmission between them, warranting further investigation into the modulatory roles of VGLUT2-ENs on extrinsic afferents.

To the best of our knowledge, our study is the first to systematically trace the anatomy of VGLUT2-ENs. We employed AAV1-cre for sparse labeling of VGLUT2-ENs through focal injection into the colon wall. The relatively low transfection efficiency of AAV1, combined with the sparse presence of VGLUT2-ENs in myenteric ganglia, allowed us to trace 31 neurons in the colorectum. The remaining 44 labeled VGLUT2-ENs could not be distinctly traced due to interference from other labeled neurons in the field of view, limiting our anatomical analysis to soma size and dendrite morphologies. For tracing, we utilized membrane-bound Channelrhodopsin2-EYFP (ChR2-EYFP), known for its effective tracing capability in previous studies to label unmyelinated visceral afferent endings more than 20 mm away from the somata in the DRG (12, 42). In our current research, ChR2-EYFP offered clearer tracing of fine dendritic structures in VGLUT2-EN somata compared to tdTomato, which fills neural somata but does not reveal fine neural processes. Our anatomical analysis revealed that most VGLUT2-ENs exhibit characteristics of Dogiel type I neurons, with small somata, lamellar-shaped dendrites, and a single long axon projecting aborally (ranging from 4 to 10 mm). Interestingly, the axonal projection length of VGLUT2-ENs appears to be significantly longer than that of motor neurons, i.e., ∼1.4 mm from a prior report with myotomy/myectomy studies (31). Only 2 out of 34 VGLUT2-ENs showed a Dogiel type II-like morphology with multiple circumferentially projecting axons. However, they are unlikely to be intrinsic primary afferent neurons (IPANs) due to their small soma size; IPANs typically have large somata (14, 26).

Our study also involves a detailed examination of the molecular profiles of VGLUT2-ENs through single-cell mRNA sequencing, using a deep read per cell (5 million). Given the rarity of VGLUT2-ENs (comprising only 1.88% of myenteric neurons), we employed fluorescence-assisted manual cell picking rather than automated cell sorting by previous studies on intestinal cells (7, 24). This manual process selectively isolated neuronal cells labeled by VGLUT2-Cre, which were confirmed to express high levels of VGLUT2 and neuronal markers while displaying low expression of the glial marker GFAP. The majority of VGLUT2-ENs express molecules essential for synaptic glutamatergic and cholinergic transmission, indicating their potential role in neural signaling. However, typical markers for descending interneurons are not prominently expressed in VGLUT2-ENs, including calcium binding proteins (calb1, calb2), CGRP (calca1, calca2), nitric oxide synthases (nos1), vasoactive intestinal peptide (vip), and serotonin synthesis/reuptake (dcc, slc6a4) (36). Further, VGLUT2-ENs show almost no expression of the mechanosensitive ion channel Piezo2, along with significantly less Piezo1 expression compared to non-VGLUT2-ENs. Functionally, interneurons are mechanosensitive and respond directly to circumferential stretch (36). The anatomical evidence mentioned above already excludes VGLUT2-ENs from the IPAN and motor neuron classes. Additionally, the evidence from single-cell RNA sequencing does not align with the classification of VGLUT2-ENs as interneurons. Therefore, our combined anatomical and molecular evidence strongly suggests that VGLUT2-ENs constitute a unique group of enteric neurons that may not directly participate in the canonical myenteric neural circuitry responsible for intestinal peristalsis and motility.

While the mechanosensitivity of enteric neurons has been extensively studied using electrophysiological methods or voltage-sensitive dye detection, these studies typically involve either dissociated, cultured preparations (21, 22), dissected neuron-muscle strip preparations with isometric stretching (38), or intact ganglia subjected to distension through intra-ganglion fluid infusion or punctate probing (23). Notably, prior research has not recorded enteric neurons while preserving the integrity of colorectal tissue and simulating stretch conditions that correlate with physiological colorectal distension. In our study, we addressed this gap by conducting optical GCaMP6f recordings from individual VGLUT2-ENs within colorectal tissues that retained intact ganglia and wall layers. We applied ascending circumferential colorectal stretch corresponding to intraluminal pressures from 0 to approximately 80-100 mmHg, well into the range of noxious distending pressure (25). To mitigate motion artifacts resulting from colorectal stretch, we introduced a novel correction method that involved subtracting the local background fluorescence from the GCaMP6f signal. This innovative correction technique enabled us to detect the onset and magnitude of evoked calcium transients with precision. The effectiveness of our correction method was validated, as we observed a nearly constant GCaMP6f signal during stretch when neural responses were pharmacologically suppressed. Our GCaMP6f recordings yield significant findings: the majority of VGLUT2-ENs, identified through focal electrical stimulation, exhibit positive responses to mechanical colorectal stretch. Furthermore, we observe that the activation of VGLUT2-EN neurons in response to colorectal stretch requires synaptic cholinergic transmission, as nicotinic cholinergic antagonist hexamethonium inhibits the responses in most VGLUT2-EN neurons. These functional results from our GCaMP6f studies on VGLUT2-ENs align remarkably well with the findings from the above scRNA-seq analysis, which indicate a low expression level of mechanosensitive ion channels (Piezo1 and Piezo2) in VGLUT2-ENs.

In conclusion, we systematically conducted anatomical, molecular, and functional studies on myenteric neurons genetically labeled by the VGLUT2 promoter using the cre-LoxP strategy. VGLUT2-ENs exhibited an even distribution throughout the colorectum, present in nearly all myenteric ganglia. Along the longitudinal axis, VGLUT2-EN density remains fairly consistent in the colonic region, with a slight decrease toward the rectal region due to the lower total number of myenteric neurons in the rectum. Interestingly, a higher proportion of VGLUT2-ENs was observed in the intermediate region between the colon and rectum, coinciding with the entry point of extrinsic afferent innervations. Tracing the anatomy of 31 VGLUT2-ENs revealed that they are predominantly Dogiel type I neurons with single aborally projecting axons ranging from 4 to 10 mm in length. Molecular profiling of VGLUT2-ENs through single-cell mRNA sequencing unveiled a unique expression profile, distinguishing them from typical interneurons. Additionally, VGLUT2-ENs displayed minimal expression of mechanosensitive ion channels (Piezo1 and Piezo2), setting them apart from mechanosensitive interneurons. Functional studies on VGLUT2-ENs under physiological colorectal distension conditions demonstrated positive responses to mechanical stretch, with nicotinic cholinergic transmission playing a pivotal role. This study provides novel insights into the distribution, anatomy, molecular profile, and function of VGLUT2-ENs, underscoring their uniqueness among enteric neurons and suggesting a potential role in modulating extrinsic afferents during colorectal function.

## Acknowledgments

This work was supported by NINDS NS113873 and NIDDK DK120824 grants awarded to Dr. Feng.

## References

1. Arshadi C, Günther U, Eddison M, Harrington KI, and Ferreira TA. SNT: a unifying toolbox for quantification of neuronal anatomy. Nature methods 18: 374–377, 2021.

2. Brehmer A. Classification of human enteric neurons. Histochemistry and Cell Biology 156: 95–108, 2021.

3. Brumovsky PR, Robinson DR, La JH, Seroogy KB, Lundgren KH, Albers KM, Kiyatkin ME, Seal RP, Edwards RH, and Watanabe M. Expression of vesicular glutamate transporters type 1 and 2 in sensory and autonomic neurons innervating the mouse colorectum. Journal of Comparative Neurology 519: 3346–3366, 2011.

4. Brumovsky PR, Robinson DR, La JH, Seroogy KB, Lundgren KH, Albers KM, Kiyatkin ME, Seal RP, Edwards RH, Watanabe M, Hokfelt T, and Gebhart GF. Expression of vesicular glutamate transporters type 1 and 2 in sensory and autonomic neurons innervating the mouse colorectum. JComp Neurol 519: 3346–3366, 2011.

5. Costa M, Spencer NJ, and Brookes SJH. The role of enteric inhibitory neurons in intestinal motility. Auton Neurosci 235: 102854, 2021.

6. Dickson EJ, Heredia DJ, McCann CJ, Hennig GW, and Smith TK. The mechanisms underlying the generation of the colonic migrating motor complex in both wild-type and nNOS knockout mice. American journal of physiology Gastrointestinal and liver physiology 298: G222–232, 2010.

7. Drokhlyansky E, Smillie CS, Van Wittenberghe N, Ericsson M, Griffin GK, Eraslan G, Dionne D, Cuoco MS, Goder-Reiser MN, and Sharova T. The human and mouse enteric nervous system at single-cell resolution. Cell 182: 1606–1622. e1623, 2020.

8. Ershov D, Phan M-S, Pylvänäinen JW, Rigaud SU, Le Blanc L, Charles-Orszag A, Conway JR, Laine RF, Roy NH, and Bonazzi D. TrackMate 7: integrating state-of-the-art segmentation algorithms into tracking pipelines. Nature methods 19: 829–832, 2022.

9. Feng B, Brumovsky PR, and Gebhart GF. Differential roles of stretch-sensitive pelvic nerve afferents innervating mouse distal colon and rectum. American journal of physiology Gastrointestinal and liver physiology 298: G402–409, 2010.

10. Feng B and Gebhart GF. Characterization of silent afferents in the pelvic and splanchnic innervations of the mouse colorectum. American journal of physiology Gastrointestinal and liver physiology 300: G170–180, 2011.

11. Feng B and Guo T. Visceral pain from colon and rectum: the mechanotransduction and biomechanics. Journal of Neural Transmission 127: 415–429, 2020.

12. Feng B, Joyce SC, and Gebhart GF. Optogenetic activation of mechanically insensitive afferents in mouse colorectum reveals chemosensitivity. American journal of physiology Gastrointestinal and liver physiology 310: G790–798, 2016.

13. Furness JB, Jones C, Nurgali K, and Clerc N. Intrinsic primary afferent neurons and nerve circuits within the intestine. Progress in Neurobiology 72: 143–164, 2004.

14. Furness JB, Kunze WA, Bertrand PP, Clerc N, and Bornstein JC. Intrinsic primary afferent neurons of the intestine. Prog Neurobiol 54: 1–18, 1998.

15. Guo T, Bian Z, Trocki K, Chen L, Zheng G, and Feng B. Optical recording reveals topological distribution of functionally classified colorectal afferent neurons in intact lumbosacral DRG. Physiological Reports 7: e14097, 2019.

16. Guo T, Patel S, Shah D, Chi L, Emadi S, Pierce DM, Han M, Brumovsky PR, and Feng B. Optical clearing reveals TNBS-induced morphological changes of VGLUT2-positive nerve fibers in mouse colorectum. American Journal of Physiology-Gastrointestinal and Liver Physiology 320: G644–G657, 2021.

17. Hockley JRF, Taylor TS, Callejo G, Wilbrey AL, Gutteridge A, Bach K, Winchester WJ, Bulmer DC, McMurray G, and Smith ESJ. Single-cell RNAseq reveals seven classes of colonic sensory neuron. Gut 68: 633–644, 2019.

18. Holton KF, Taren DL, Thomson CA, Bennett RM, and Jones KD. The effect of dietary glutamate on fibromyalgia and irritable bowel symptoms. Clin Exp Rheumatol 30: 10–17, 2012.

19. Huang Z, Liao L, Wang Z, Lu Y, Yan W, Cao H, and Tan B. An efficient approach for wholemount preparation of the myenteric plexus of rat colon. Journal of Neuroscience Methods 348: 109012, 2021.

20. Ke MT, Fujimoto S, and Imai T. SeeDB: a simple and morphology-preserving optical clearing agent for neuronal circuit reconstruction. Nat Neurosci 16: 1154–1161, 2013.

21. Kugler EM, Michel K, Kirchenbüchler D, Dreissen G, Csiszár A, Merkel R, Schemann M, and Mazzuoli-Weber G. Sensitivity to Strain and Shear Stress of Isolated Mechanosensitive Enteric Neurons. Neuroscience 372: 213–224, 2018.

22. Kugler EM, Michel K, Zeller F, Demir IE, Ceyhan GO, Schemann M, and Mazzuoli-Weber G. Mechanical stress activates neurites and somata of myenteric neurons. Frontiers in cellular neuroscience 9: 342, 2015.

23. Mazzuoli G and Schemann M. Mechanosensitive enteric neurons in the myenteric plexus of the mouse intestine. PLoS One 7: e39887, 2012.

24. Morarach K, Mikhailova A, Knoflach V, Memic F, Kumar R, Li W, Ernfors P, and Marklund U. Diversification of molecularly defined myenteric neuron classes revealed by singlecell RNA sequencing. Nature neuroscience 24: 34–46, 2021.

25. Ness TJ and Gebhart GF. Colorectal distension as a noxious visceral stimulus: physiologic and pharmacologic characterization of pseudaffective reflexes in the rat. Brain Res 450: 153–169, 1988.

26. Nurgali K, Stebbing MJ, and Furness JB. Correlation of electrophysiological and morphological characteristics of enteric neurons in the mouse colon. Journal of Comparative Neurology 468: 112–124, 2004.

27. Preibisch S, Saalfeld S, and Tomancak P. Globally optimal stitching of tiled 3D microscopic image acquisitions. Bioinformatics 25: 1463–1465, 2009.

28. Qu Z-D, Thacker M, Castelucci P, Bagyanszki M, Epstein ML, and Furness JB. Immunohistochemical analysis of neuron types in the mouse small intestine. Cell and tissue research 334: 147–161, 2008.

29. Raab M and Neuhuber WL. Glutamatergic Functions of Primary Afferent Neurons with Special Emphasis on Vagal Afferents. In: International Review of Cytology: Academic Press, 2007, p. 223–275.

30. Ren J, Hu HZ, Liu S, Xia Y, and Wood JD. Glutamate receptors in the enteric nervous system: ionotropic or metabotropic? Neurogastroenterology & Motility 12: 257–264, 2000.

31. Sang Q, Williamson S, and Young H. Projections of chemically identified myenteric neurons of the small and large intestine of the mouse. The Journal of Anatomy 190: 209–222, 1997.

32. Sang Q and Young H. Chemical coding of neurons in the myenteric plexus and external muscle of the small and large intestine of the mouse. Cell and tissue research 284: 39–53, 1996.

33. Schneider CA, Rasband WS, and Eliceiri KW. NIH Image to ImageJ: 25 years of image analysis. Nature methods 9: 671–675, 2012.

34. Siri S, Maier F, Santos S, Pierce DM, and Feng B. Load-bearing function of the colorectal submucosa and its relevance to visceral nociception elicited by mechanical stretch. American journal of physiology Gastrointestinal and liver physiology 317: G349–G358, 2019.

35. Smith-Edwards KM, Najjar SA, Edwards BS, Howard MJ, Albers KM, and Davis BM. Extrinsic Primary Afferent Neurons Link Visceral Pain to Colon Motility Through a Spinal Reflex in Mice. Gastroenterology 157: 522–536.e522, 2019.

36. Smith TK, Spencer NJ, Hennig GW, and Dickson EJ. Recent advances in enteric neurobiology: mechanosensitive interneurons. Neurogastroenterol Motil 19: 869–878, 2007.

37. Spencer NJ, Nicholas SJ, Robinson L, Kyloh M, Flack N, Brookes SJ, Zagorodnyuk VP, and Keating DJ. Mechanisms underlying distension-evoked peristalsis in guinea pig distal colon: is there a role for enterochromaffin cells? American journal of physiology Gastrointestinal and liver physiology 301: G519–527, 2011.

38. Spencer NJ and Smith TK. Mechanosensory S-neurons rather than AH-neurons appear to generate a rhythmic motor pattern in guinea - pig distal colon. The Journal of physiology 558: 577–596, 2004.

39. Swaminathan M, Hill-Yardin EL, Bornstein JC, and Foong JP. Endogenous glutamate excites myenteric calbindin neurons by activating group I metabotropic glutamate receptors in the mouse colon. Frontiers in neuroscience 13: 426, 2019.

40. Wahba G, Hebert A-E, Grynspan D, Staines W, and Schock S. A rapid and efficient method for dissociated cultures of mouse myenteric neurons. Journal of neuroscience methods 261: 110–116, 2016.

41. Wang G-D, Wang X-Y, Xia Y, and Wood JD. Dietary glutamate: interactions with the enteric nervous system. Journal of Neurogastroenterology and Motility 20: 41, 2014.

42. Zhu Y, Feng B, Schwartz ES, Gebhart GF, and Prescott SA. Novel method to assess axonal excitability using channelrhodopsin-based photoactivation. J Neurophysiol 113: 2242–2249, 2015.

